# Leptin protects against the development and expression of cocaine addiction-like behavior in heterogenous stock rats

**DOI:** 10.1101/2021.07.22.453426

**Authors:** L.L.G. Carrette, C. Coral, C. Crook, B. Boomhower, M. Brennan, C. Ortez, K. Shankar, S. Simpson, L. Maturin, G. de Guglielmo, L. Solberg Woods, A. A. Palmer, O. George

## Abstract

In addition to its pleasurable effects, weight control is a significant contributor to initiation, maintenance and relapse of cocaine use. This suggests that individual differences in bodyweight control and feeding hormones, such as leptin may contribute to the vulnerability to cocaine use disorder. While pre-clinical studies have shown a mutually inhibitory relationship between leptin and cocaine, they have used small sample sizes and did not investigate individual differences in a large heterogeneous population. Here, we tested if individual differences in bodyweight and blood leptin level is associated with high or low vulnerability to addiction-like behaviors using data from 500 heterogenous stock rats and 160 blood samples from the Cocaine Biobank, using a model of extended access to intravenous self-administration of cocaine. Finally, we tested a separate cohort to evaluate the causal effect of exogenous leptin administration on cocaine seeking. Bodyweight, while changing due to cocaine self-administration in males, was not related to the vulnerability to addiction-like behavior. Blood leptin levels after ~6 weeks of cocaine self-administration did not correlate with addiction-like behaviors, however, baseline blood leptin levels before any access to cocaine negatively predicted addiction-like behavior. Finally, administration of leptin reduced cocaine intake after acute withdrawal and cocaine seeking after 6 weeks of protracted abstinence. These results demonstrate that high blood leptin level before access to cocaine may be a protective factor against the development of cocaine addiction-like behavior, that exogenous leptin reduces the motivation to take and seek cocaine, but that blood leptin level and bodyweight changes in current users are not good biomarkers for addiction-like behaviors.

## Introduction

Cocaine is used primarily for its pleasurable effects by young adults. Overall, 5.5 million Americans reported using cocaine in 2019, which includes 1 million with cocaine use disorders (SAMHSA, 2020). While most research on cocaine is focused on its rewarding effect, there is evidence that weight control is a significant contributor to the initiation and maintenance of cocaine use, especially in females (Bruening et al., 2018; Cochrane et al., 1998). Moreover, increased weight gain during cocaine abstinence further hinders recovery and leads to an increase in relapse (Billing and Ersche, 2015). These results suggest that individual differences in bodyweight control and feeding hormones, such as leptin which mediates food satiety (Friedman, 2014; Schwartz et al., 1996), may contribute partially to cocaine use disorder vulnerability. However, the exact role of cocaine self-administration on bodyweight and leptin level and the effect of leptin on cocaine seeking is unclear, partly due to the lack of longitudinal studies and experiments with low sample size or identical inbred animals.

Converging lines of evidence suggest that cocaine may alter leptin levels and that leptin may modulate cocaine addiction-like behaviors. As both target the dopaminergic system, an interaction between self-administration of cocaine and food rewards has been observed (Carroll et al., 1989; Figlewicz et al., 2003; Fulton et al., 2006; Hommel et al., 2006). Pre-clinical studies have shown that cocaine has acute and transient anorexigenic effects (Balopole et al., 1979), followed by a compensatory increase in the consumption of fat and carbohydrates without significant weight gain (Bane et al., 1993) associated with increased metabolism (Billing and Ersche, 2015) and reduced leptin levels (You et al., 2019). On the other hand, leptin signaling attenuates the effects of cocaine addiction-like behavior in rats (Lee et al., 2018; Shen et al., 2016; You et al., 2019). Similar results on the effect of cocaine on feeding have been found in clinical studies (Castro et al., 1987; Ersche et al., 2013; Weddington et al., 1990), but the relationship with leptin is less clear. Reduced leptin levels were reported for crack cocaine users (Ersche et al., 2013; Escobar et al., 2018; Levandowski et al., 2013), but a controlled cocaine-administration study found no effect on leptin levels (Bouhlal et al., 2017). Moreover, opposite to the results in rats, increased leptin levels were reported to lead to higher cocaine craving (Martinotti et al., 2017), leaving the role of variable leptin levels on cocaine seeking unclear.

To address this gap in the literature we tested the hypothesis that cocaine self-administration leads to an increased bodyweight in abstinence through reduced blood leptin levels that contribute to increased cocaine seeking in outbred heterogeneous stock (HS) rats (Baud et al., 2013; Hansen and Spuhler, 1984) that mimic the diversity of the human population (Woods and Mott, 2017). A sub-hypothesis was that leptin may be a protective factor against the development of cocaine addiction-like behavior. To this end, we used an animal model of extended access (12 h per day for ~3 weeks) to intravenous cocaine self-administration with evaluation of cocaine intake (fixed ratio), motivation to seek cocaine (progressive ratio), resistance to punishment (contingent footshock), and irritability-like behavior (bottle brush test). At the end of the behavioral phenotyping, rats were assigned an addiction index (AI), which is a composite Z-score of the different addiction-like behaviors and separated into groups with high addiction (HA) or low addiction (LA)-like behaviors through a median split. To evaluate the impact of cocaine self-administration on bodyweight and blood leptin levels, we used phenotypic data from just under 500 animals and 160 blood samples form 100 rats from the Cocaine Biobank (Carrette et al., 2021). We also used a cohort of rats to test the causal effect of exogenous leptin administration on cocaine seeking.

## Methods

### Animals

Male and female RFID tagged HS rats (Rat Genome Database NMcwiWFsm #13673907, sometimes referred to as N/NIH) were provided by Dr. Leah Solberg Woods. Rats were shipped at 3-4 weeks of age, quarantined and then pair-housed on a 12 h/12 h normal light/dark cycle (lights off at 9:00 AM) in a temperature (20-22°C) and humidity (45-55%) controlled vivarium with ad libitum access to tap water and food pellets (PJ Noyes Company, Lancaster, NH, USA). All of the procedures were conducted in strict adherence to the National Institutes of Health Guide for the Care and Use of Laboratory Animals and were approved by the Institutional Animal Care and Use Committees of The Scripps Research Institute and UC San Diego. Experiments were performed in cohorts of 48-60 rats. In a first experiment, we used behavioral data from 470 animals from the Cocaine Biobank (Carrette et al., 2021). Next, we used 30 males and 29 females to test the effect of leptin on cocaine seeking. At completion of the experiments (18 weeks after catheter surgery) 19 rats either died of unknown cause or had failed catheter as tested using a short acting anesthetic (brevital) and were excluded from the study, leaving 23 males and 17 females. Finally, behavioral data and blood samples from 30 males and 30 females at two timepoints (baseline and withdrawal) were used from the biobank.

### Drugs

Cocaine HCl (National Institute on Drug Abuse, Bethesda, MD) was dissolved in 0.9% sterile saline and administered intravenously at a dose of 0.5 mg/kg/infusion. Volume of infusion is 100 ul. Leptin was dissolved in 0.9% sterile saline and administered intravenously at a dose of 0.6 mg/kg/infusion.

### Jugular vein catheterization

This surgery inserts a catheter into the jugular vein to allow for intravenous cocaine self-administration. Rats were anesthetized with vaporized Isoflurane (1-5%). Intravenous catheters were aseptically inserted into the right jugular vein using the procedure described previously (Carrette et al., 2021). Catheters consisted of Micro-Renathane tubing (18 cm, 0.023-inch inner diameter, 0.037-inch outer diameter; Braintree Scientific, Braintree, MA, USA) attached to a 90 degree angle bend guide cannula (Plastics One, Roanoke, VA, USA), embedded in dental acrylic, and anchored with mesh (1 mm thick, 2 cm diameter). Tubing was inserted into the vein following a needle puncture (22G) and secured with a suture. The guide cannula was punctured through a small incision on the back. The outside part of the cannula was closed off with a plastic seal and metal cover cap, which allowed for sterility and protection of the catheter base. Flunixin (2.5 mg/kg, *s.c.*) was administered as analgesic, and cefazolin (330 mg/kg, i.m.) as antibiotic. Rats were allowed three days for recovery prior to any self-administration. They were monitored and flushed daily with heparinized saline (10 U/ml of heparin sodium; American Pharmaceutical Partners, Schaumberg, IL, USA) in 0.9% bacteriostatic sodium chloride (Hospira, Lake Forest, IL, USA) that contained 52.4 mg/0.2 ml of cefazolin.

### Behavioral phenotyping

The rats were subjected to only one behavioral procedure per day.

#### Operant self-administration

The rats were allowed to self-administer cocaine individually in operant chambers (29 cm × 24 cm × 19.5 cm; Med Associates, St. Albans, VT, USA) that were enclosed in lit, sound-attenuating, ventilated environmental cubicles and computer controlled. Each chamber was equipped with two retractable levers that would only be extended for the duration of the test. Pressing of the right, active lever activated an infusion pump on a fixed ratio (FR1) schedule to deliver the drug infusion through plastic catheter tubing over 5 seconds, followed by a 20 time out during which pressing the lever had no scheduled consequences. Right lever presses also activated a cue light that remained illuminated during the time-out. Pressing the left, inactive lever had no scheduled consequences. The rats went through 10 short access (ShA) sessions of 2 h and 14 long access (LgA) sessions of 6 h. Five sessions were performed per week, with a break over the weekend.

*Progressive Ratio (PR) responding* was tested to assess motivation. Number of lever presses required to receive a drug infusion increase progressively (according to the following schedule: 1, 2, 4, 6, 9, 12, 15, 20, 25, 32, 40, 50, 62, 77, 95, 118, 145, 178, …). When the rat did not achieve the required number of presses for the next infusion in the hour following the last infusion, the session was terminated with the breakpoint defined as the last achieved ratio. PR tests were performed after ShA, after LgA and after foot shock tests. For one cohort, 2 additional PR tests were performed with intravenous (*i.v.*) leptin treatment (0.6 mg/kg in saline) or saline vehicle, 30 min before the session using a Latin square design.

#### Compulsive-like responding using contingent foot shock

In this self-administration session, 30% of active lever presses were randomly associated with a foot shock (0.3 mA for 0.5 s), delivered through the floor grid.

#### Bottle brush test for irritability

Rats were placed in the back of a clean cage and approached by a rotating bottle brush for 5 s, The brush was then slowly withdrawn while rotating for 5 s. The test was repeated 10 times with 10 s intervals between each test. Three observers independently scored defensive irritability-like behaviors: escape, jumping, vocalization, grooming, and digging; and aggressive irritability-like behaviors: exploration, boxing, biting, following, and latency biting. Baseline behavior was tested before the first SA session and compared to the levels after LgA (18h in withdrawal).

#### Reinstatement (RI) responding

The subset of animals that were tested for cocaine seeking with leptin treatment were re-introduced in the SA chambers for 2 h with access to the levers after 6 weeks of forced abstinence. The right lever press still activated a cue light, but did not result in drug delivery. Three sessions were preformed, separated by 1 week of abstinence. Session 1 and 3 animals received *i.v.* saline, session 2 animals received *i.v.* leptin (0.6 mg/kg in saline), 30 minutes before the start of the session.

### Blood collection and analysis

#### Baseline blood (before ShA)

Retroorbital bleeds were performed under anesthesia with the eye numbed by a topical ophthalmic anesthetic (proparacaine hydrochloride).

#### Terminal

Blood was collected through cardiac puncture after CO_2_ inhalation. Each cohort was divided into three final endpoints. Intoxication: euthanized with drug on board after a 3h SA session. Withdrawal: euthanized 18h in withdrawal after a LgA session. Abstinence: euthanized ~3-4 weeks after the last cocaine intake during SA sessions.

Fresh blood was collected in EDTA coated tubes to avoid coagulation and was centrifuged at 2000 g at room temperature (RT) for 10 min. The supernatant plasma was immediately transferred per 500 µl aliquots into fresh tubes, scored for quality on a scale from 0-6, snap frozen on dry ice and stored at −80°C. Frozen aliquots were thawed on ice and analyzed using a Leptin Rat ELISA kit (Invitrogen).

### Fat pad collection

For the one cohort of rats that received leptin treatment, fat pads were collected at sacrifice. For female rats, retroperitoneal and parametrial fat was collected and weighed. For male rats, retroperitoneal and epididymis fat was collected and weighed. The length of the animals (with and without tail) was also measured to calculate body mass index (BMI).

### Data Analysis

Data was analyzed with Graphpad Prism and RStudio. Statistical tests were performed as described. Post hoc tests were Tukey corrected.

- An *addiction index (AI)* for each rat was calculated as described before (Carrette et al., 2021) using the average of the Z-scores for intake, motivation, withdrawal, and compulsivity. Escalation index (intake): E = Z_f_ / Z_0_, where Z_f_ is the Z score of an animal on the last 3 days of escalation and Z_0_ is the Z score of an animal during the first day of escalation. A daily escalation index (Ei) can be calculated for each session (E_i_ = Z_i_ / Z_0_) where Z_i_ is the Z-score on a given escalation session i. This index can also be calculated for the 1st hour or the entire session. Irritability index (withdrawal): Z-score of the total irritability score with the baseline scores subtracted. PR index (motivation): Z-score of the breakpoint after LgA. Shock index (compulsivity): Z-score of the breakpoint.

- *Body mass index (BMI):* (bodyweight / body length^2) × 10

## Results

### Bodyweight differences in abstinence from cocaine compared to naive

The timeline for the testing of cocaine addiction-like behavior is illustrated in **Figure 1A**. Animals are equipped with an intravenous (*i.v.*) catheter for cocaine self-administration (SA) during 10 short (ShA) and 14 long (LgA) sessions. This gives them the opportunity to escalate their intake and develop dependency. Withdrawal is tested by comparing their irritability 18 h in withdrawal to the baseline level before SA. Compulsivity is tested using 1 session with shock as an aversive consequence to lever pressing and motivation is tested after ShA, LgA and shock. Given the genetic diversity of the animals, a wide variability in addiction-like behaviors between the animals can be observed. To better characterize the animals, Z-scores of these behaviors were calculated (**Figure 1B**) and averaged in an overall addiction index (AI) per animal. Animals with a positive AI were classified as having high addiction-like behaviors (HA), those with a negative AI as having low addiction-like behaviors (LA) Mixed 3-way ANOVA revealed that the 2 addiction groups were significantly different from each other (p group < 0.0001). When looking at the bodyweight of the animals at the 3 different timepoints for sacrifice (intoxicated, in withdrawal immediately following cocaine self-administration, or in protracted abstinence 1 month later, see Figure 1A), the weights in abstinence are significantly higher than 4-5 weeks earlier in withdrawal and intoxication in male rats (post hoc test following 3-way ANOVA, p < 0.0001). (**Figure 1C**). Moreover, when comparing with the naive littermates, for males we see an effect of addiction-like behavior on bodyweight. In abstinence, the HA group has a significantly lower bodyweight than the naive littermates (post hoc test following 3-way ANOVA: p < 0.001). In females, while not significant, there is the same trend for bodyweight increase during abstinence compared to intoxication and withdrawal, but there is an opposite trend for weight gain in HA compared to naive littermates.

**Figure 1.**
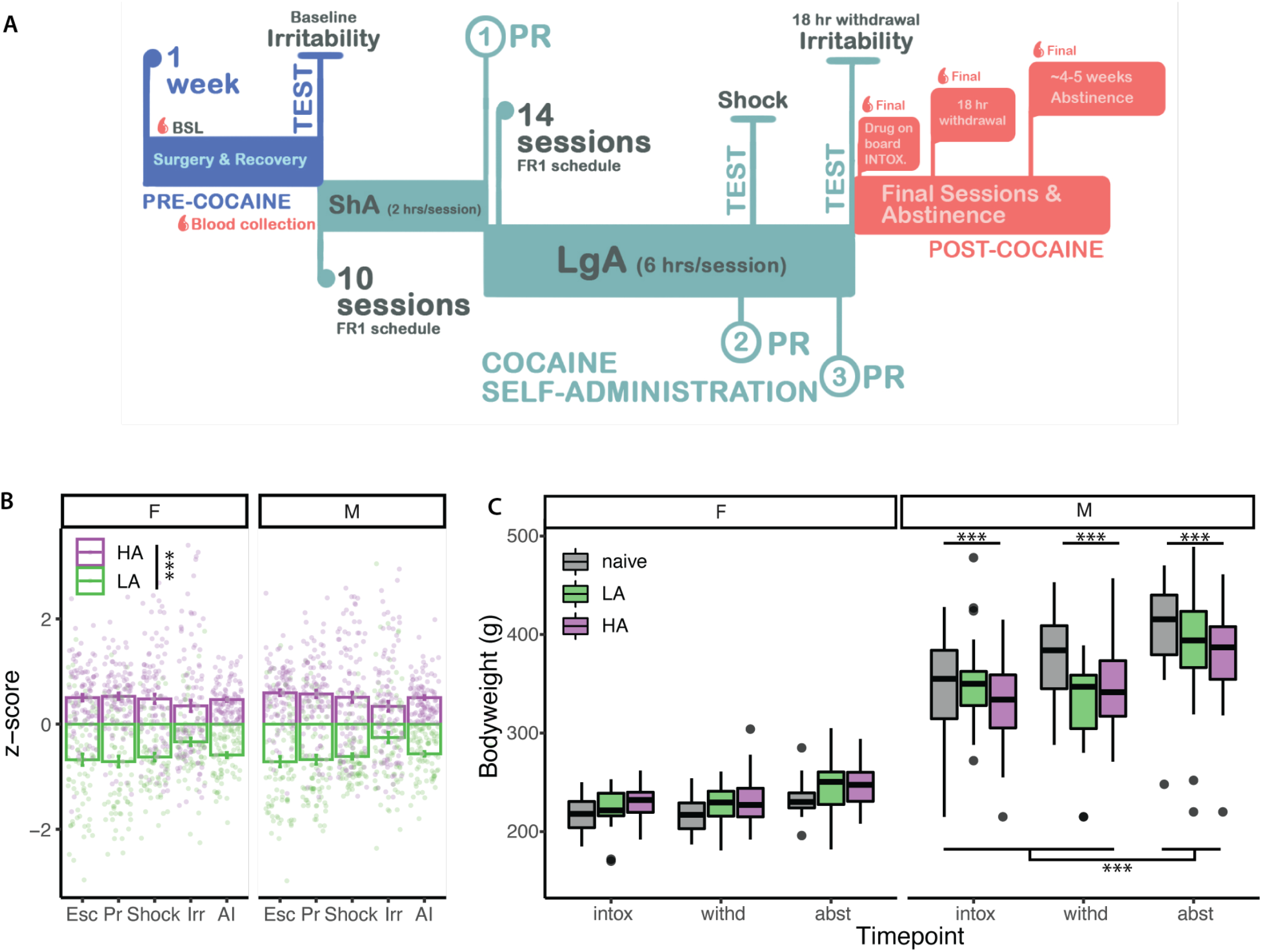
Bodyweights of HS rats. **(A)** Cocaine self-administration behavioral timeline. **(B)** Individual escalation (Esc), motivation (Pr), compulsivity (Shock), withdrawal (Irr) Z-scores and addiction index (AI) show the distinction of the animals with high (HA) and low (LA) addiction-like behaviors. Mixed 3-way ANOVA with score as within, sex and group as between variables: p group < 0.0001, p sex = 0.44, p score = 0.34, p group:sex = 0.84, p group:score < 0.0001, p sex:score = 1, p group:sex:score = 0.90) (N = 185 F + 193 M). **(C)** Bodyweight at one of 3 timepoints of HA and LA animals compared to naives shows opposite trends in females and males (3-way ANOVA, p timepoint < 0.0001, p sex < 0.0001, p group < 0.5, p timepoint:sex < 0.0001, p timepoint:group = 0.32, p sex:group < 0.01, p timepoint:sex:group = 0.15. Post hoc: p naive M-HA M < 0.001 and p abst M – intox/withd M < 0.0001) (HA: N = 106 F + 100 M, LA: N = 79 F + 93 M, naive: N = 46 F + 46 M, equally distributed over the timepoints).

### Systemic leptin infusion reduces cocaine seeking

Given the role leptin has as a mediator of bodyweight, we next decided to investigate this further in a cohort of HS animals (N = 30 F + 30 M) after the SA protocol. **Figure 2A** shows the timeline of the experiments added to the standard behavioral protocol of Figure 1A for this cohort. First, two additional PR experiments were added in which the leptin was administered *i.v*. in a Latin square design immediately following SA, to test the effect of leptin on the motivation for cocaine (**Figure 2B**). Both male and female rats significantly reduced their cocaine intake after leptin treatment (mixed 2-way ANOVA: p treatment = 0.017, p sex = 0.12, p treatment:sex = 0.85). Alternatively, the animals can be divided by their addiction-like behaviors. Per definition of the groups, there is a significant difference between the groups in intake (p group < 0.0001) and leptin reduces intake similarly in both groups (mixed 2-way ANOVA, p treatment = 0.0036, p treatment:group = 0.74). There are too few animals for 3-way ANOVA combining sex and addiction group. The animals were then kept in their home cage for 6 weeks of forced abstinence. Since the bodyweight differences became most pronounced during abstinence, 12 h of food intake during the dark cycle was measured every 2 weeks of abstinence to see if a difference in food intake caused the weight differences (**Figure 2C**). Food intake normalized by bodyweight was significantly different between the sexes (p sex < 0.0001) and slightly decreased over time (p time = 0.045), without interaction (p time:sex = 0.080). There was no difference in food intake between the 2 addiction groups (mixed 2-way ANOVA, p time = 0.18, p group = 0.52, p time:group = 0.40). Next, the effect of leptin on cocaine seeking in abstinence in the absence of cocaine was assessed by 3 reinstatement sessions over 3 weeks. During these sessions the animals were returned to the SA boxes, where they could press for but did not receive cocaine. Due to extinction, the effect of reinstatement decreases over time. This was somewhat mitigated by 1 week of forced abstinence in between the 3 sessions. In week 1, reinstatement was tested with vehicle treatment. Week 2 tested the effect of leptin treatment. Finally, vehicle treatment was repeated in week 3 to confirm that the effect was due to leptin and not extinction. Both male and female rats responded to the treatment with leptin significantly reducing lever presses compared to both control reinstatement sessions (mixed 2-way ANOVA, p treatment = 0.003, p sex = 0.04, p treatment:sex = 0.31) (**Figure 2D**). When comparing the reinstatement of HA and LA animals, there was an interaction between groups and treatment, with leptin having a significant effect on HA rats (post hoc test p < 0.001) and not the LA, which reinstate significantly less than the HA group (mixed 2-way ANOVA, treatment p = 0.0068, group p = 0.0077, interaction p = 0.0070). After the experiments, while still in abstinence, the animals were weighed and sacrificed. There was only a significant difference between the sexes for bodyweight (2-way ANOVA, p sex < 0.0001, p group = 0.12, p sex:group = 0.89) (**Figure 2E**). The same non-significant results for fat pad weight and BMI were obtained (data not shown). Endogenous leptin levels were determined in blood plasma (data not shown). Here too, the only significant difference was between males and females (2-way ANOVA, p sex < 0.0001, p group = 0.21, p interaction = 0.23). While the levels were identical in females, a small trend could be observed with HA males having slightly reduced leptin levels. Correlating weight-normalized plasma leptin levels with the addiction index reached no significance (M: R^2^ = 0.08, p = 0.18; F: R^2^ = 0.009, p = 0.72; all: R^2^ = 0.04, p = 0.20) (**Figure 2F**). Usually, leptin levels positively correlate with bodyweight, but **Figure 2G** shows that this correlation is lost in abstinence, especially in the females (M: R^2^ = 0.12, p = 0.11; F: R^2^ = 0.05, p = 0.38).

**Figure 2.**
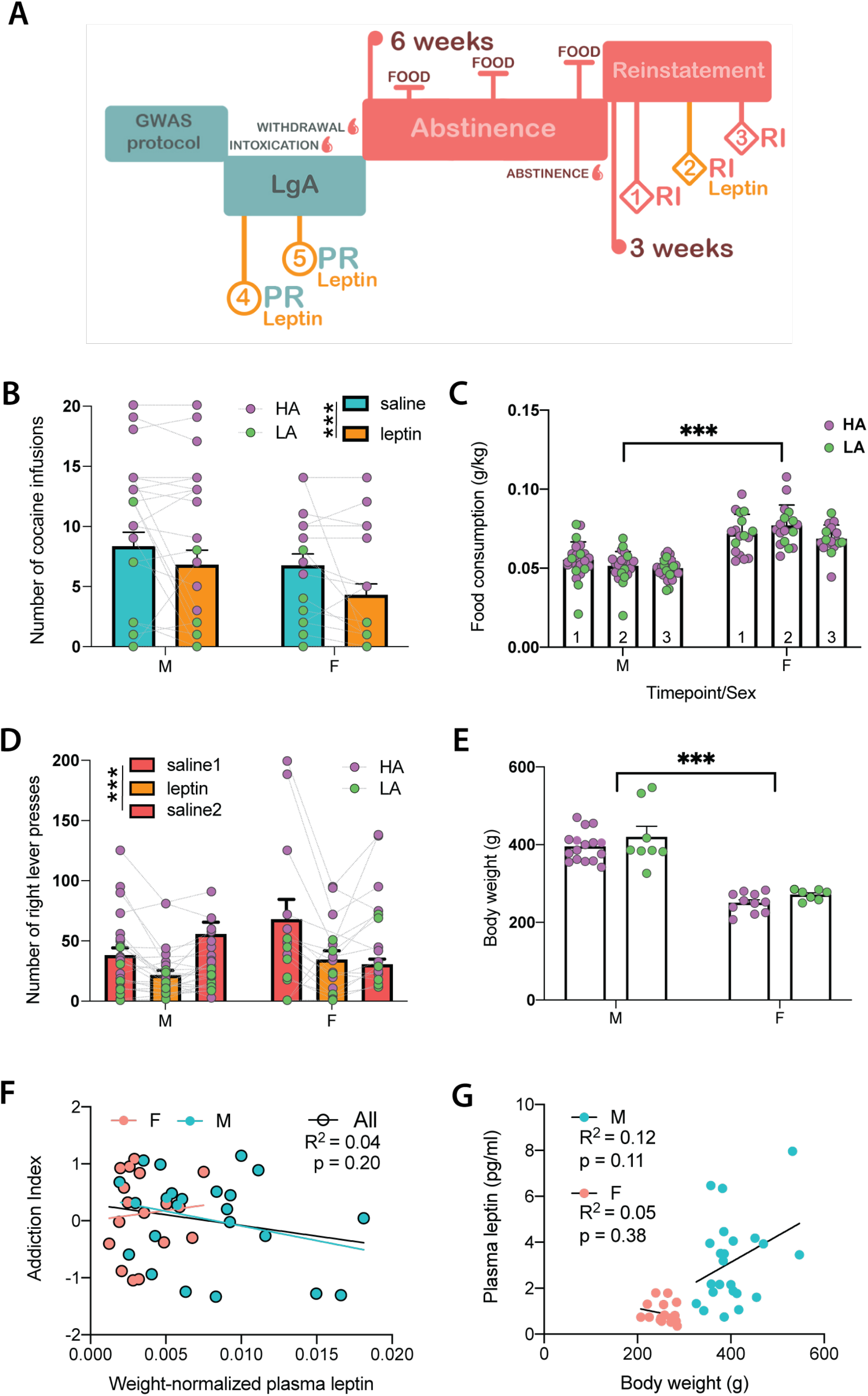
Additional behavior, weight, and leptin testing in one cohort of animals. **(A)** Timeline of the additional experiments for better characterizing the effect of exogenous leptin on cocaine seeking, and the effect of cocaine abstinence on food intake and weight. **(B)** Both males and female rats significantly reduced their cocaine intake after leptin treatment (mixed 2-way ANOVA: p treatment = 0.017, p sex = 0.12, p treatment:sex = 0.85). (**C**) There was no difference in food intake between the 2 addiction groups (HA/LA) (mixed 2-way ANOVA: p time = 0.18, p group = 0.52, p time:group = 0.40). **(D)** Both males and females pressed significantly less during reinstatement after leptin treatment (mixed 2-way ANOVA, p treatment = 0.003, p sex = 0.04, p treatment:sex = 0.31). **(E)** Males weigh significantly more than females, without a difference between addiction groups (2-way ANOVA: p sex < 0.0001, p group = 0.12, p sex:group = 0.89). **(F)** The addiction index of the animals did not significantly correlate with their weight-normalized plasma leptin levels, in males, females or combined (M: R2 = 0.08, p = 0.18; F: R2 = 0.009, p = 0.72; all: R2 = 0.04, p = 0.20). **(G)** The plasma leptin levels in abstinence did not correlate with the bodyweight in both male and female rats (M: R2 = 0.12, p = 0.11; F: R2 = 0.05, p = 0.38).

### Endogenous leptin level at baseline protects against the development of addiction-like behaviors

To circumvent the influence of variable cocaine intake on leptin levels, next leptin was analyzed in plasma samples from HS rats at baseline out of the cocaine biobank (N = 30 F + 30 M) (Carrette et al., 2021), before exposure to cocaine. At baseline, the plasma leptin levels correlate with bodyweight for both sexes (**Figure 3A**) (M: R^2^ = 0.29, p = 0.026; F: R^2^ = 0.16, p = 0.029). The weight-normalized leptin levels further correlate with the AI that these animals end up developing later in life (**Figure 3B**) (M: R^2^ = 0.15, p = 0.04; F: R^2^ = 0.11, p = 0.086; all: R^2^ = 0.13, p = 0.006). The correlation is weak, and lost when the sexes are split, but higher endogenous levels of leptin seem to provide some protection against the development of more severe addiction-like behaviors.

**Figure 3.**
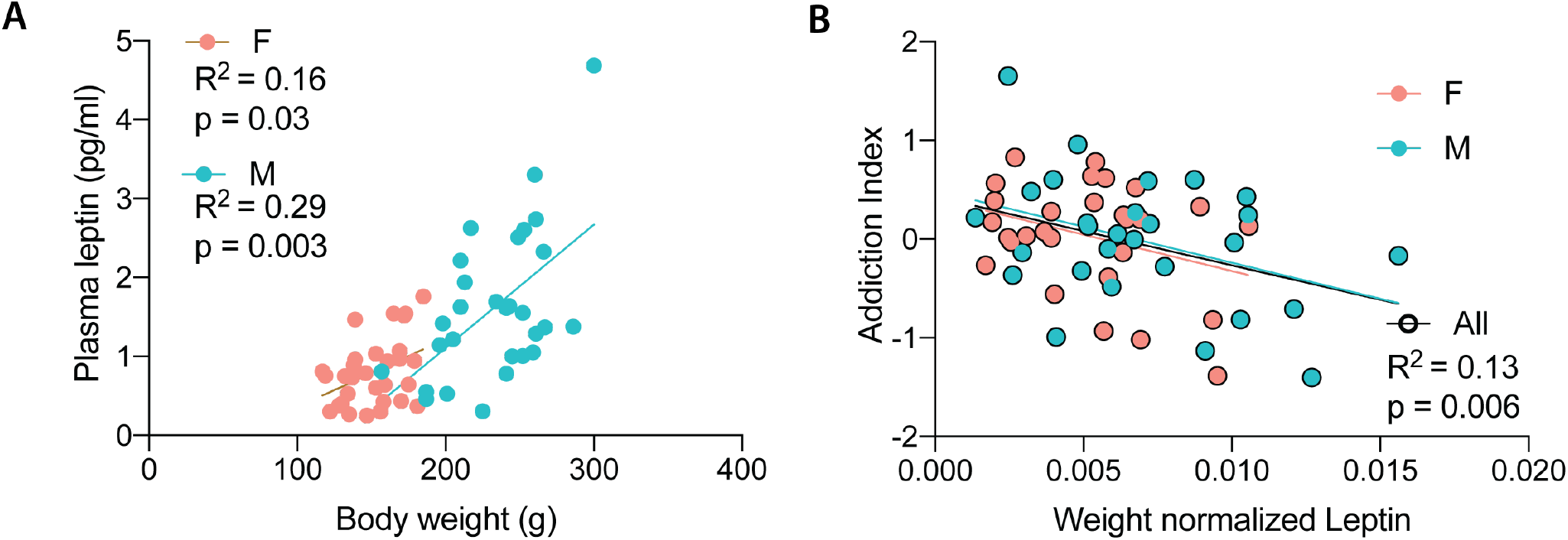
Plasma leptin analysis at baseline. **(A)** Baseline leptin levels positively correlate to bodyweight (M: R2 = 0.29, p = 0.026; F: R2 = 0.16, p = 0.029). **(B)** AI negatively correlate with weight normalized baseline leptin levels (M: R2 = 0.15, p = 0.04; F: R2 = 0.11, p = 0.086; all: R2 = 0.13, p = 0.006).

## DISCUSSION

Here we examined the effect of cocaine self-administration on bodyweight and leptin levels, and the effect of leptin on cocaine seeking in diverse HS rats. Contrary to our working hypothesis, male HA animals showed a significant lower bodyweight compared to naive controls. Abstinent HA females showed a trend towards a higher bodyweight in abstinence, but this did not reach significance. There was no relation between the bodyweight and addiction-like behaviors. Plasma leptin levels in abstinence seemed dysregulated by cocaine intake as they lost their typical positive correlation with bodyweight, but no consistent differences in HA vs LA animals were found. However, when looking at leptin levels in these animals before the first cocaine self-administration, a correlation was found with their future AI. This indicates that endogenous leptin may be a protective factor against the development of cocaine addiction-like behavior. This effect was further validated by suppression of cocaine seeking in progressive ratio during withdrawal and reinstatement during abstinence upon leptin administration.

Leptin is the best-known mediator of bodyweight, through modulating food intake and food metabolism both homeostatically in the hypothalamic circuitry (Farooqi et al., 1999; Halaas et al., 1995) and by reducing dopamine (DA) signaling in relation to food cues in the mesolimbic circuitry (Hommel et al., 2006; Sadaf Farooqi et al., 2007; Van Der Plasse et al., 2015). It has been reported to interact with cocaine cues and intake. In rats, leptin injections, systemically or intracranial in the VTA or NAc have been shown to attenuate cocaine conditioned place preference (CPP), cocaine seeking under extinction conditions and cocaine induced locomotor activity (Lee et al., 2018; You et al., 2016). In mice, both the injection of the leptin receptor antagonist or deletion of the receptor was shown to enhance cocaine CPP (Shen et al., 2016). We extended these findings to show that exogenous leptin inhibits cocaine self-administration under a progressive ratio and showed that the effect on seeking was maintained 6 weeks in abstinence with reinstatement tests. No sex difference was found in this effect, which could have been masked by the high dose of leptin and require a response curve with lower doses. When looking at the effect of LA vs HA animals, we found that while leptin suppressed PR in both groups, only the reinstatement of HA animals got affected by the leptin treatment.

Additionally, endogenous leptin levels are altered by cocaine. In human, the increase (Levandowski et al., 2013; Martinotti et al., 2017), decrease (Escobar et al., 2018), and a decreasing trend (Ersche et al., 2013) or lack of effect (Bouhlal et al., 2017) have been described, that could be resulting from the different study protocols and addiction time points. In rats, it was shown that cocaine intake and even the expectancy of intake decreases leptin levels, which normalize in abstinence (You et al., 2016). The trend of lower leptin levels in HA males that was observed here could be a remainder of that suppression. The lack of significant findings, might be due to the loss of animals in this long (18-week) experiment and with it statistical power, as we were also unable to detect significant differences in bodyweight, fat pad weight, BMI, food intake and leptin levels between the groups (except for sex differences). This reveals in any case that leptin and bodyweight are likely bad biomarkers for addiction-like behavior.

The Cocaine Biobank (Carrette et al., 2021) is a unique resource providing phenotypic data on a large collection of HS rats who underwent state-of-the-art extended access to intravenous cocaine self-administration self-administering cocaine, which is highly relevant to cocaine use disorder (Edwards and Koob, 2013; George et al., 2014). As a result of their diverse genotypes, HS rats exhibit diverse addiction-like behavior phenotypes (HA vs LA), which mimic diversity in a human population (Woods and Mott, 2017), but under the strictly controlled conditions of a pre-clinical study that are hard to achieve in human subjects. Using baseline samples from the cocaine biobank, we were uniquely able to look at leptin levels before cocaine exposure and correlate them with future addiction-like behaviors. This revealed a correlation of baseline plasma leptin levels with the addiction index these animals will develop. While the predictive effect is small, it is in line with the protective effect of leptin treatment we observed against cocaine seeking, potentially related to the effect of leptin mutations in mood disorders (Atmaca et al., 2008, 2002; Kraus et al., 2001), and emphasizes a crucial role of leptin in the brain reward system. Finally, it illustrates the presence of the many subtle differences in a diverse population that contribute to individual vulnerability to addiction-like disorders.

In conclusion, leptin and cocaine have a mutual interactive effect. While leptin and bodyweight seemed affected by cocaine, we were unable to show a clear effect that relates with addiction-like behaviors. On the other hand, we did show that increased exogenous and endogenous leptin levels reduce cocaine reward and protect individuals from developing substance use disorders. The latter could help explain individual differences in the propensity to develop cocaine use disorder and might hold potential for the development of the first treatment against cocaine use disorder.

## Acknowledgements

We thank the Preclinical Addiction Research Consortium. The research was supported by NIDA (U01DA043799 and U01DA043799). OP and AAP were also supported by P50DA037844 and DA043799.

